# Unraveling micro-architectural modulations in neural tissue upon ischemia by Correlation Tensor MRI

**DOI:** 10.1101/2021.02.20.432088

**Authors:** Rita Alves, Rafael Neto Henriques, Leevi Kerkelä, Cristina Chavarrías, Sune N Jespersen, Noam Shemesh

**Affiliations:** Champalimaud Research, Champalimaud Centre for the Unknown, Lisbon Portugal; UCL Great Ormond Street Institute of Child Health, University College London, London, United Kingdom; Center of Functionally Integrative Neuroscience (CFIN) and MINDLab, Clinical Institute, Aarhus University, Aarhus, Denmark; Department of Physics and Astronomy, Aarhus University, Aarhus, Denmark

## Abstract

Noninvasively detecting and characterizing modulations in cellular scale micro-architecture is a *desideratum* for contemporary neuroimaging. Diffusion MRI (dMRI) has become the mainstay methodology for probing microstructure, and, in ischemia, its contrasts have revolutionized stroke management. However, the biological underpinnings of the contrasts observed in conventional dMRI in general and in ischemia in particular are still highly debated since the markers only indirectly reporter on microstructure. Here, we present Correlation Tensor MRI (CTI), a method that rather than measuring diffusion, harnesses diffusion correlations as its source of contrast. We show that CTI can resolve the sources of diffusional kurtosis, which in turn, provide dramatically enhanced specificity and sensitivity towards ischemia. In particular, the sensitivity towards ischemia nearly doubles, both in grey matter (GM) and white matter (WM), and unique signatures for neurite beading, cell swelling, and edema are inferred from CTI. The enhanced sensitivity and specificity endowed by CTI bodes well for future applications in biomedicine, basic neuroscience, and in the clinic.

## INTRODUCTION

Progressive modulation in tissue micro-architecture is associated with diverse natural neural processes including development^1^, plasticity^2^, memory^3^, learning^4,5^, connectivity between brain areas^6,7^, ageing^8^, and recovery from injury^9^. Adverse micro-architectural alterations in the neural tissue milieu are also associated with psychiatric disorders such as depression^10^, neurodegenerative diseases such as Parkinson’s disease^11^ and Alzheimer’s disease^12^, and injuries such as ischemic stroke^9^ and traumatic brain injury^13^. In ischemic stroke – one of the leading causes of disability and death worldwide^14^ – a complex cascade of micro-architectural events^15^ occurs acutely following a blood vessel occlusion and the ensuing metabolic and aerobic deprivation. These micro-architectural modifications include neurite beading^16,17^, intracellular swelling due to loss of ion homeostasis, cytotoxic edema, and cell death (Fig. 1a) followed later by disruptions in the blood-brain barrier, vasogenic edema and tissue clearance. The extent of these processes later determines the prognosis, the potential for functional recovery^18^ and success of treatment.

**Figure 1.**
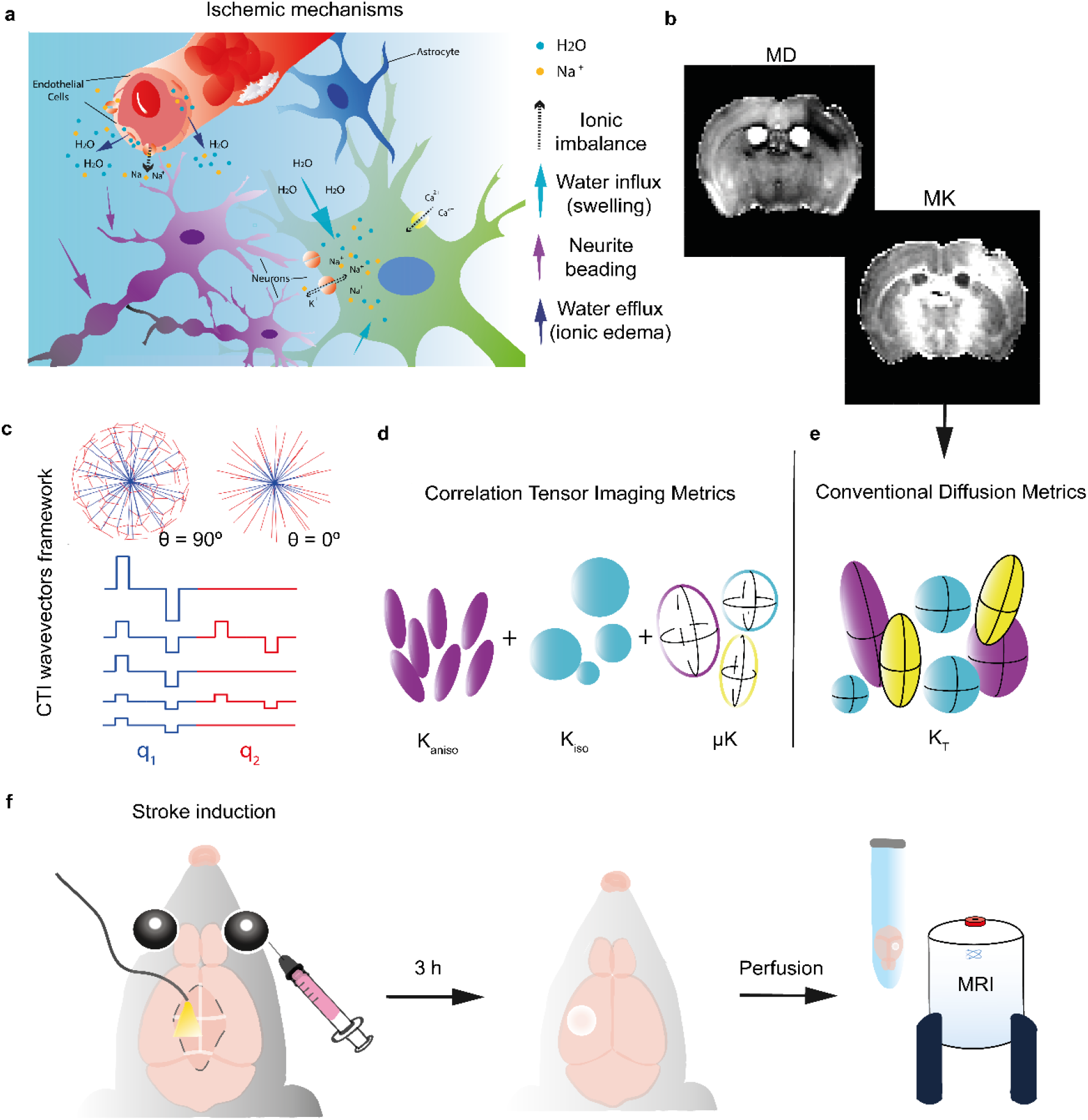
Acute ischemic mechanisms, CTI elements illustration and stroke experimental set up. **(a)** At an acute ischemic phase – after vascular occlusion – microscopic disturbances such as ionic imbalance (dashed), cell swelling (light blue arrows account for an excessive cellular influx of water in endothelial and neuronal cells), with a subsequent neurite beading (purple) are occurring (cytotoxic edema). This event is followed by ionic edema – transcapillary flux of Na^+^ and water efflux (dark blue arrows). The blood brain barrier remains intact at this stage. **(b)** MD and total kurtosis (MK) are presented as **(e)** conventional diffusion metrics. Although conventional diffusion metrics obtained from single diffusion encoding (SDE) techniques assess quantitative information on tissue heterogeneity, these cannot resolve the different kurtosis sources, hence lacking specificity and hampering an accurate association with biophysical alterations. On the other hand, correlation tensor imaging (CTI), relying on the cumulant expansion of Double Diffusion Encoding (DDE) signal with **(c)** 90° and 0° combinations of wavevectors, enables the disentanglement of the kurtosis sources **(d)** (Anisotropic Kurtosis, K_aniso_; Isotropic Kurtosis, K_iso_; and Microscopic Kurtosis, µK), providing a more specific characterization of the microscopic alterations occurring in ischemic tissue. **(f)** A photothrombotic model was used in order to induce a well delimited focal infarct. Upon longitudinal incision and target region coordinates identification (S1bf), irradiation on the left hemisphere was performed for 15 minutes. At 3 h post irradiation offset, the brains were fixed via transcardial perfusion, extracted and placed in an NMR tube with Fluorinert for MRI imaging.

Given the progressive nature of micro-architectural modulations in neural tissue, non-invasive mapping of tissue microstructure^19–22^ plays a pivotal role in assessing enhancements in microstructure e.g., due to learning, as well as adverse effects upon disease or neural injury. Diffusion-weighted magnetic resonance imaging (dMRI) has become the mainstay in contemporary neuroimaging for probing tissue microstructure. The dMRI methodology non-invasively imparts contrast sensitive to diffusion-driven displacement of water molecules in the tissue^23^, thereby indirectly probing their interaction with the microscopic boundaries imposed by the cellular environment^21^. In stroke, dMRI provided the first effective means of early stroke detection^24,25^, thereby facilitating the administration of treatment within the limited time window of operation. Effectively, dMRI is sensitive to the displacement distribution induced by the microstructural features in the tissue. The displacement distribution can be characterized by an apparent diffusion coefficient (ADC) along a single direction. Diffusion Tensor MRI (DTI^26^) has been developed to estimate the 3-dimensional diffusion tensor. However, the diffusion tensor represents an approximation of the displacement distribution to a 3D normal distribution, and therefore it can only capture Gaussian diffusion. Diffusion Kurtosis MRI (DKI^27^), was thus introduced to quantify the amount of non-Gaussian diffusion in the tissue, thereby providing increased sensitivity towards microstructure^28^. Effectively, DTI quantifies the mean of the distribution of apparent displacement distribution and DKI estimates the non-Gaussianity of the distribution. Indeed, dMRI has completely revolutionized our ability to follow microstructural modulations over extended periods of time^29^.

However, the non-Gaussianity of the displacement distribution can arise from numerous microstructural sources, which are all conflated in conventional dMRI. The main limitation stems from the use of “Single Diffusion Encoding” (SDE^30,31^): acquisitions imparting sensitivity towards molecular displacement in a single epoch for a specific single orientation. In such a scenario mesoscopic effects such as orientation dispersion of anisotropic microenvironments, the size distribution of microscopic restricted systems, and the microscopic kurtosis itself (arising from structural disorder or restricted diffusion) – all potential consequences of microstructural modulations – are conflated and cannot be resolved without extensive assumptions. The SDE signal thus does not carry sufficient information for distinguishing different microstructural features and resolving these contributions remains elusive. In stroke, for example, effects arising from cross-sectional variations due to neurite beading^16,32^ (Fig. 1a, purple), cell body swelling^33^ (Fig. 1a, light blue) or the transcapillary ionic edema^34^ (Fig. 1a, dark blue), are notoriously difficult to decipher. Notably, the biophysical underpinnings of even the simplest ADC changes observed in stroke^24,35^ have been vigorously debated for the last three decades^16,36,37^. The parameters estimated from such SDE approaches, including mean diffusivity (MD), fractional anisotropy (FA), or total kurtosis^38^ (K_T_) thus suffer from lack of specificity.

Here, we depart from the convention of mapping the diffusion and kurtosis tensors using SDE, and rather quantitatively map the *displacement correlation tensor*^39^ using Double Diffusion Encoding – a method applying two diffusion-sensitizing epochs that probe the correlation between diffusion-driven displacements in different directions^31,40,41^. The ensuing Correlation Tensor MRI (CTI^42^) approach facilitates the disentanglement of the different underlying non-Gaussian diffusion sources, namely: anisotropic kurtosis^43^, K_aniso_; isotropic kurtosis^44^, K_iso_; and microscopic kurtosis^32,45^, μK (Fig.1d). In particular, K_aniso_ quantifies the degree of anisotropy in the tissue irrespective of orientation dispersion effects; K_iso_ quantifies the variance in the sizes of the microscopic diffusion environments; and μK quantifies the degree of non-Gaussian effects induced by structural disorder^32,46^ and restricted diffusion^45^. Each of these sources can represent a fundamentally different property of the underlying microstructure, and therefore their measurement can potentially greatly enhance specificity. We present the first CTI experiments in a stroke model, which evidence unique signatures for the different underlying micro-architectural modulations, including beading, swelling and edema. Importantly, the sensitivity towards stroke detection is dramatically increased by this approach. CTI thus delivers the sought-after specificity and sensitivity towards microstructural features, which can impact our understanding the progressive modulations occurring in tissue in health and disease.

## RESULTS

### Focal thrombi induction

Upon injection with photosensitive dye and irradiation with the appropriate wavelength (Fig.1f) for generation of reactive oxygen species, that in turn produce severe endothelial damage and thrombi formation, a well-delineated extensive stroke was evidenced in T_2_ weighted images (Fig. 2a) 3h post-ischemia. The affected area appeared with strong hyperintense contrast, unilaterally covering the barrel cortex and to some extent the hippocampus. The dMRI signals averaged across all acquired directions (powder-averaged^41^) also clearly delineated the stroke area as hyperintense signals (Fig. 2b). For all five mice that underwent ischemic induction, the stroke area was highly reproducible (Supplementary Fig. S1). By contrast, no interhemispheric differences could be observed in the control mice that were injected with the photosensitive dye but did not undergo irradiation (Fig. 2c-d). Both T2-weighted images and powder-averaged diffusion-weighted images appear symmetric and without abnormal contrasts.

**Figure 2.**
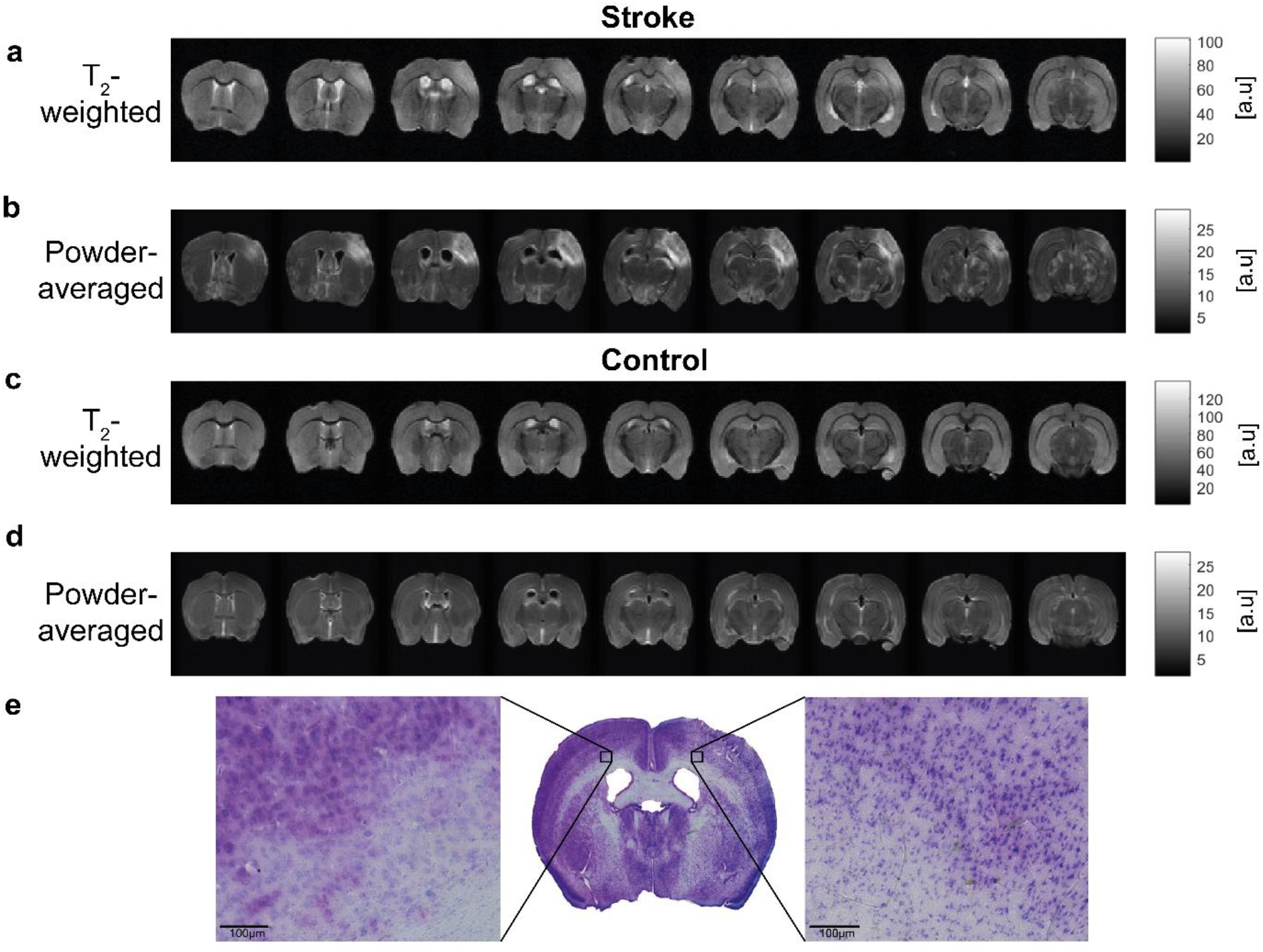
Raw T2-weighted, powder-averaged data and histological validation of the lesion. **(a, c)** T2-weighted images and **(b, d)** the powder-averaged signal decays computed by averaging the diffusion-weighted signals decays over 135 directions of diffusion wave vectors from representative extracted brains from both stroke and control groups (total b-value = 3000 s/mm^2^). 9 representative coronal slices are presented from rostral to caudal direction. The T2-weighted contrast shows elevated intensity values around the infarcted region (left hemisphere, barrel cortex) GM, whereas the powder-averaged contrast presents elevated values for both GM and WM within the infarcted region. **(e)** Histological analysis was performed to assess cell damage and validate the lesion. A cresyl violet stained brain section is presented. By enabling the labelling of Nissl corpuscles present in neurons, it shows the lesion in the subcortical region of the left hemisphere from a representative mouse brain perfused at 3 hours post ischemic stroke. A critical selected GM region in the ipsilesional hemisphere was magnified (53x), presenting pale staining, associated with micro vacuolation of the cytoplasm, and less intact cells associated with cell damage, when compared to the intact contralesional region (left).

A histological evaluation of the stroked area (Fig. 2e) clearly demonstrated abnormalities in Nissl (Cresyl violet) staining. The zoomed in view of the ipsilesional cortex (Fig. 2e, right) also shows the reduced staining and density, compared with the contralesional cortex (Fig. 2e, left).

### Conventional contrast in stroke

#### Total kurtosis and Mean diffusivity

As in prior studies^47,48^, we observed strongly elevated total kurtosis values in the stroked group. Fig. 3a shows the clearly elevated values of total kurtosis (K_T_) in a representative mouse brain in the affected hemisphere, as well as reduced MD and slightly reduced FA (Supplementary Fig. S2). The K_T_ values appeared elevated in both GM and WM, and the contrast was more apparent compared with the MD contrast. On the other hand, no interhemispheric differences were observed in K_T_ in the control group (c.f. Fig. 3b for a representative control brain) and other diffusion tensor metrics (MD, FA were also symmetric (Supplementary Fig. S2). However, it is difficult to draw conclusions on the sources for stroke-induced changes in K_T_ or MD which could involve any of the mechanisms described in Fig.1a, as also represented in the illustration of kurtosis sources (Fig. 3c).

**Figure 3.**
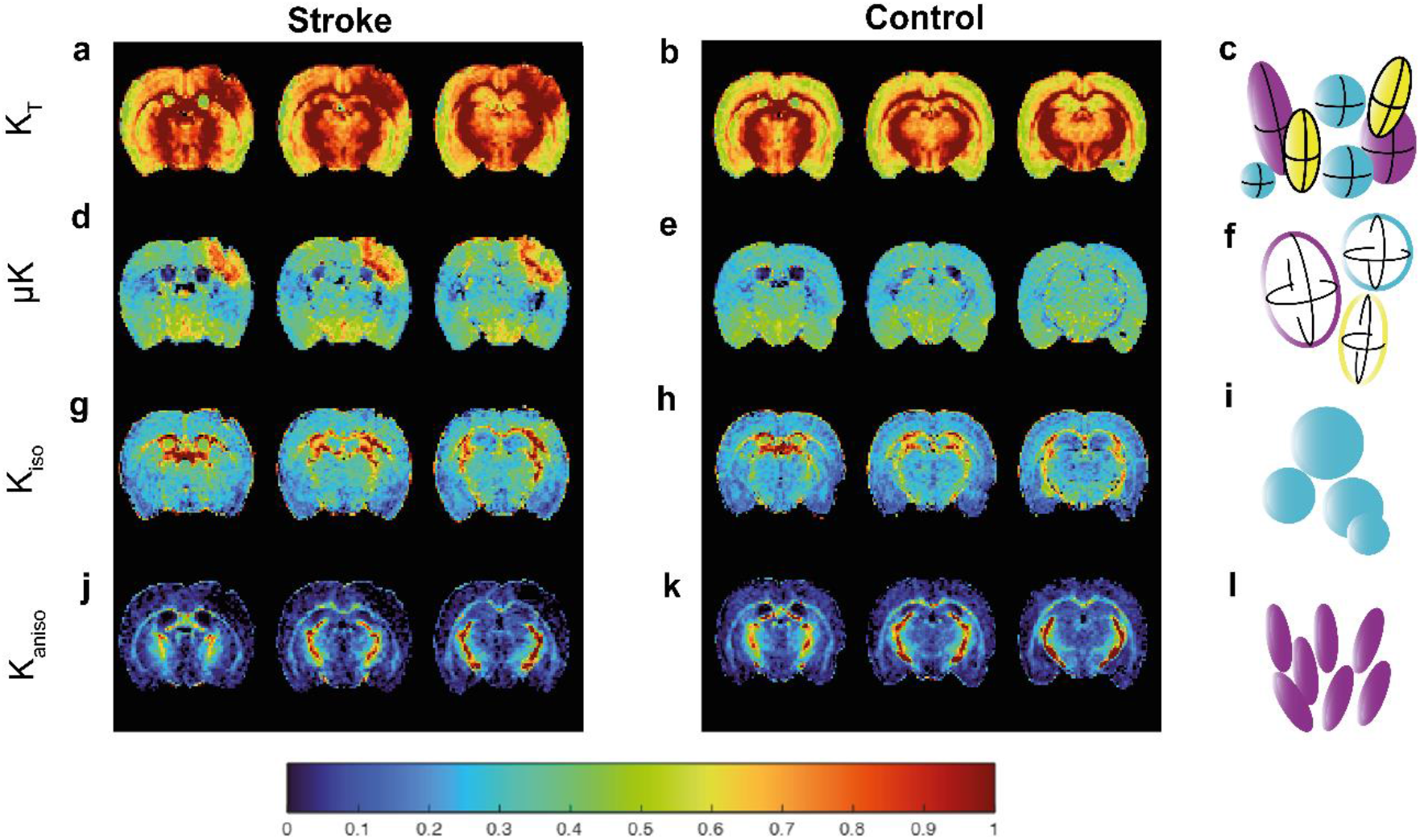
Kurtosis sources maps for stroked and control brains. Kurtosis maps (**(a, b)** K_T_, **(d, e)** μK, **(g, h)** K_iso_ and **(j, k)** K_aniso_) for three slices from representative brains from the stroke (left) and control (right) groups are presented with respective illustration of each kurtosis source **(c, f, i, l)**. In the ischemic group, the ipsilesional hemisphere shows higher K_T_ intensity values in GM and a clear distinction between hemispheres is observed in μK values, showing greater intensities in both WM and GM. K_aniso_ presents lower values in both WM and GM within the ipsilesional hemisphere when compared to the contralesional hemisphere.

### CTI contrast in stroke

Given that CTI can further disentangle the different sources, we next turn to evaluate the differences observed in CTI’s metrics.

#### Microscopic Kurtosis source (μK)

The μK metric derived from the CTI analysis in the stroked brains – which represents contributions from structural disorder and restricted diffusion to non-Gaussian diffusion – was dramatically elevated in the affected area (Fig. 3d). This observation was consistent for all N = 5 mice (Supplementary Fig. S3), and a visual examination shows that the elevated μK appeared higher both in GM and WM tissues. It is further noteworthy that the stroke was much better delineated in the μK maps compared with the conventional total kurtosis counterparts (c.f. below for quantitative analyses). In the control group, again the μK maps showed no striking interhemispheric differences (Fig. 3e), suggesting that the elevated μK in stroke, representing structural disorder / restriction (Fig. 3f), is not due to some asymmetric left-right (or other) imaging artifact.

#### Isotropic kurtosis source (K_iso_)

The isotropic kurtosis contrast, which represents the variance in diffusion tensor traces in the tissue derived from CTI analysis exhibited increases, mainly evident in WM, and much less noticeable contrast differences in GM (Fig. 3g). No apparent differences in WM or GM tissues were observed between hemispheres for K_iso_ in the control group (Fig. 3h), suggesting that the variance in isotropic tensor magnitudes (Fig. 3i) in the stroked group is again not related with some imaging source.

#### Anisotropic kurtosis source (K_aniso_)

The anisotropic kurtosis (K_aniso_) contrast in stroke, which is proportional to the degree of intravoxel anisotropy irrespective of orientation dispersion – evidenced strong reductions, more evident in the ipsilesional GM (Fig.3j) but also showing some reductions in WM. In the control group, the same K_aniso_ contrast did not appear different between the hemispheres (Fig.3k). In other words, the GM in the stroked hemisphere was characterized by decreased local anisotropy, independent of local orientation (Fig.3l).

To examine whether the stroked area could be better delineated using CTI metrics we plotted a 3D render of the μK maps and compared them to their conventional total kurtosis counterpart in a representative brain (Fig. 4). The entire 3D render can be viewed in Supplementary Videos, and the rendered data can be viewed for every animal in Supplementary Fig. S4. These depictions of the stroke reveal how μK provides a much more sensitive assessment of the affected area. Both the area affected, and the magnitude of the effect are much more obvious in the μK 3D rendering (Fig. 4).

**Figure 4.**
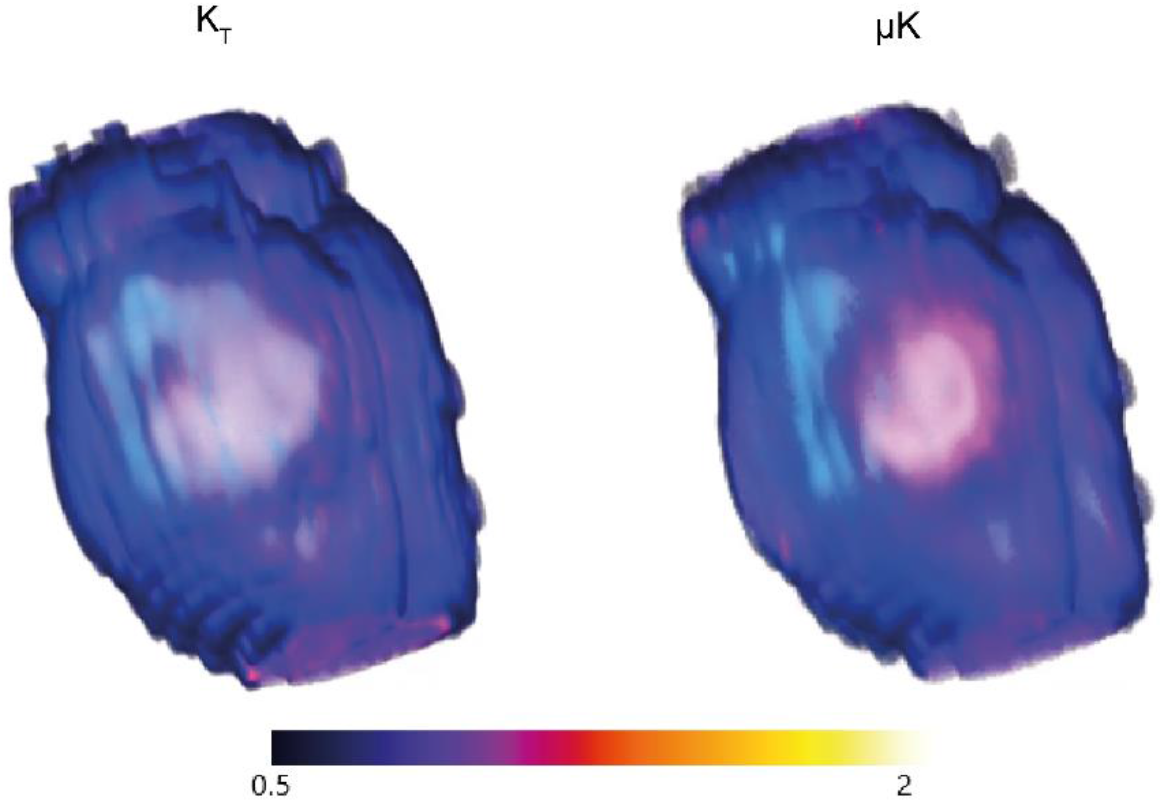
3D rendering maps for K_T_ and µK. Volume render maps using tricubic smooth interpolation for K_T_ and μK sources in a representative stroked group brain (Supplementary Videos and Supplementary Fig. S4). μK shows a more elevated contrast in the ipsilesional hemisphere (delineating the region affected region by stroke) in comparison to K_T_.

To assess whether the CTI metrics provide a more sensitive evaluation of stroke, we analysed the percent changes for all affected voxels (GM and WM combined (Supplementary Fig. S6) as well as for GM and WM separately. Fig. 5a shows the magnitude of the effects (with respect to the contralesional hemisphere in the stroked group. While total kurtosis changes ∼40% of its nominal value in stroke, the μK contrast in GM+WM was nearly double, with ∼80% change of its nominal values. While K_T_ changes were quite similar between GM and WM, the μK changed more dramatically in WM (∼120%) and still very strongly in GM (∼70%). K_iso_ on the other hand, changed only by ∼40% in general, with higher affinity to WM (∼70% change but with large standard error). Finally, K_aniso_ decreased by over 70% overall, with nearly ∼-80% changes in GM and ∼-50% changes in WM.

**Figure 5.**
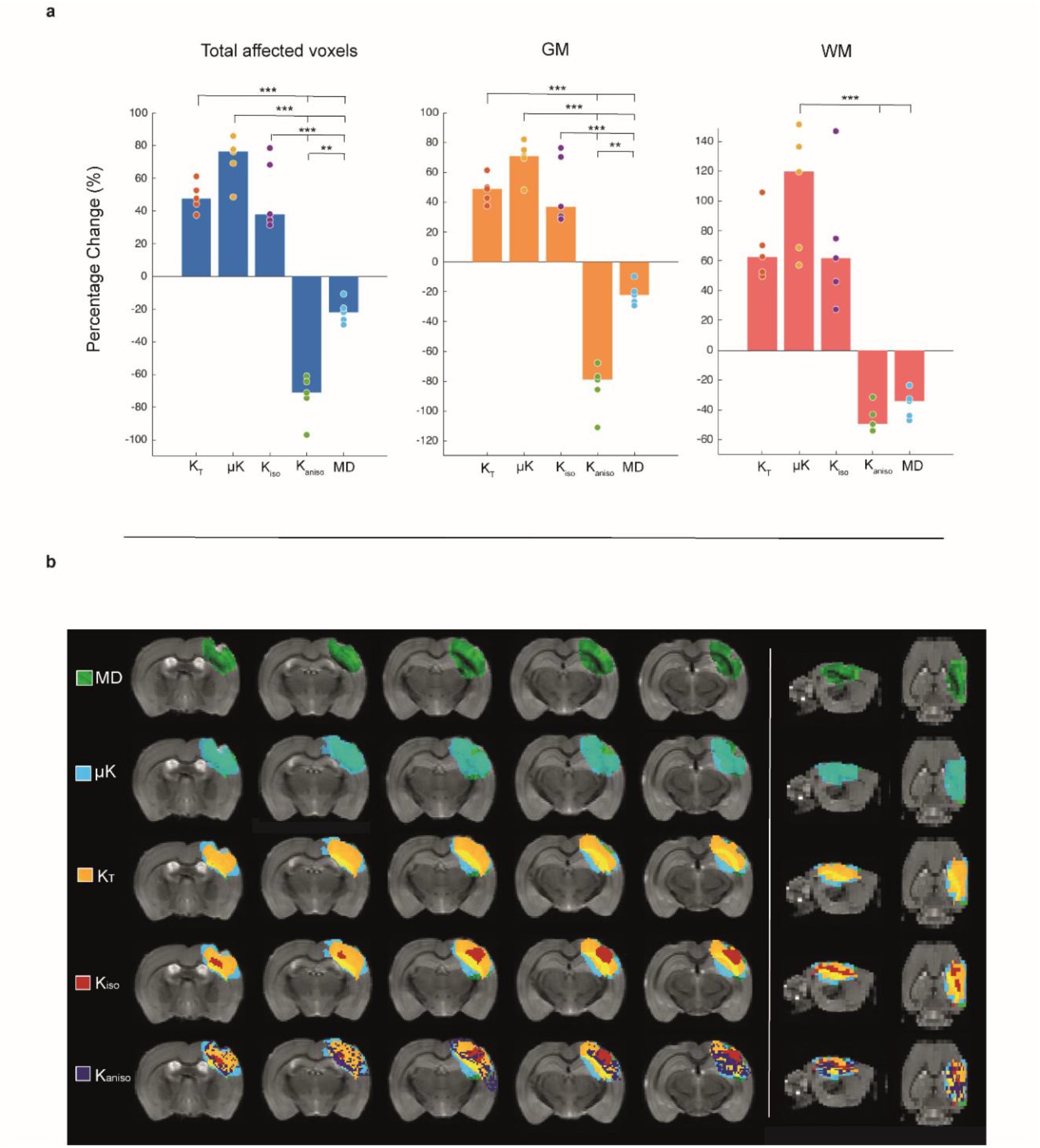
Interhemispheric percent change and ipsilesional affected voxels overlays. **(a)** Ipsilesional-contralesional ratios accounting for the mean of all voxels for different ROIs – GM and WM – from five stroked brains (N = 5) were assessed for K_T_, μK, K_iso_, K_aniso_ and MD. Median values are presented for total affected voxels, GM and WM. Percentage change was distinctly more elevated for both GM and WM in μK. Statistically significant differences (p < 0.001) were reported between the ratios of K_T_, μK, K_iso_ and K_aniso_ and MD in both GM and WM; **(b)** Metrics addition analysis using MRIcroGL in b_0_ averaged images of a representative stroked brain. 2D renderings of overlay ROI maps on the ipsilesional hemisphere were performed according to the voxels affected by the lesion in each metric. Metrics ROIs were overlayed by the following order: MD (green), μK (cyan), K_T_ (yellow), K_iso_ (red) and K_aniso_ (dark purple). Coronal slices are presented, showing that μK highlights a greater area than MD for affected voxels, followed by K_T_, K_iso_ and K_aniso_. On the right, the sagittal (left hemisphere) and axial views presenting the lesioned region are displayed following the same order of ROI and estimates overlay.

We then assessed the overlap of CTI metrics with their more conventional counterparts by overlaying the different areas found to be affected by each metric (Fig. 5b). The ensuing overlay analysis clearly demonstrates that μK detects more affected voxels overall, including in the lesion periphery, with a higher affinity towards GM; areas with large overlap between the sources indicates multitude microstructural effects, while areas characterized solely by µK represent more subtle effects on the tissue. Areas with large K_iso_ and µK overlap appear more confined to the core of the stroke. Supplementary Fig. S6 shows that these observations are highly consistent within other animals.

### *In vivo* CTI in stroke

To ensure that the effects above were not dominated by fixation effects upon perfusion of the brains, and to show the *in vivo* feasibility of the approach we repeated the same experiments at the 3h time point post-ischemia in an *in vivo* setting. Supplementary Fig. S7 shows that the same trends as observed in the *ex vivo* results reported above persist; most importantly, the dramatic elevation of μK persists in all tissues, K_iso_ is increased in WM, and K_aniso_ decreases in GM. Therefore, all the trends above are reproduced *in vivo*.

## DISCUSSION

Alterations in neural tissue micro-architectural features accompany a multitude of biological processes ranging from learning, development and plasticity to aging, neurodegeneration and neural injury. Therefore, longitudinally and noninvasively characterizing such time-dependent processes has been a *desideratum* for contemporary neuroimaging. Indeed, diffusion-driven methods such as DWI, DTI and DKI have played a crucial role in stroke evaluation owing to the microscopic-scale sensitivity arising from water diffusion within cellular-scale barriers. For example, the diffusion coefficient (or tensor) based methods can detect the stroke at very early stages^24^, while the DKI counterparts show larger lesion sizes^48^. However, these methods suffer from a degeneracy that prevents them from being specific towards microstructural features.

Here, the CTI methodology was shown to impart both sensitivity and specificity towards microscopic modulations within the imaged voxels upon ischemia. By relying on the measurement of the 4^th^ order displacement correlation tensor using DDE, CTI disentangles the features contributing to non-Gaussian diffusion into the underlying sources: anisotropic, isotropic and microscopic kurtosis sources (while also providing the more conventional metrics typically used for stroke imaging (e.g., MD, K_T_, FA). Indeed, the CTI approach dramatically improved the detection and characterization of the stroked region. In particular, (some of) its metrics provided a much more sensitive detection of the ischemic features, while the combination of its parameters provided enhanced specificity and insights into the underlying micro-architectural modulations in the tissue.

For instance, μK showed a dramatic increase and much higher sensitivity to the ischemia compared to its conventional K_T_ counterpart. The μK maps revealed both much stronger contrast than the conventional metrics as well as larger areas affected by the stroke. The reason for this enhanced sensitivity can be ascribed to the positive (μK, K_iso_) and negative (K_aniso_) effects that negate each other in K_T_, but not in μK, which reflects the microstructure without the confounds of local anisotropy effects. Therefore, μK is a strong candidate for earlier detection of ischemia and better assessment of potential stroke severity, functional outcomes and prognosis.

From a more biological perspective, the subacute phase of ischemia investigated in this study is characterized by three major micro-architectural modulations occurring downstream of the disruption in oxygen supply. In particular (1) neurite beading^16^ due to neuronal dysfunction and preceding cell death^49^; and (2) edema formation^34^ (more free water in the tissue) and disruption of extracellular/intracellular ionic balance; and (3) cell swelling^33^ (enlargement of cellular structures). It is thus instructive to assess how these processes could affect the CTI metrics.

Figure 6 shows simulations for different micro-architectural scenarios. The upper panel describes the different CTI metrics for different degrees of beading. Interestingly, we find clear signatures for increased beading in CTI signals. The MD decreases (consistent with Budde et al.^16^), while total kurtosis increases, which can now be explained by large μK increases accompanied by small decreases in K_aniso_ (K_iso_ remains zero for all beaded scenarios by definition^42^). This, importantly, contrasts with the scenario of edema formation (c.f. Fig.6b), which shows that as the edema fraction increases (from only beads to only free water, 0 < f_edema_ < 1), μK and K_aniso_ decrease in a similar fashion, while MD increases and total kurtosis decreases. In this case, K_iso_ first increases as the diffusivity difference between the beads and the more freely diffusing water is initially large but as the free fraction begins to dominate, the variance in diffusivities is strongly skewed to free diffusion, culminating in totally free diffusion, which has zero variance (K_iso_ = 0). Finally, if we consider edema formation in the presence of “swelling” of disconnected spherical objects (e.g., representing spheres, Fig. 6c), K_aniso_ remains constant and zero while K_T_ and K_iso_ evidence maxima in their values, and μK is negative, and increasing in value as edema fraction increases. Hence, CTI provides unique insights into the underlying modes of micro-architectural modulations and edema formation. While MD decreases can occur due to changes in diffusivities only, the combination of MD decreases, µK increases, and decreasing K_aniso_ can be considered hallmarks of beading.

**Figure 6.**
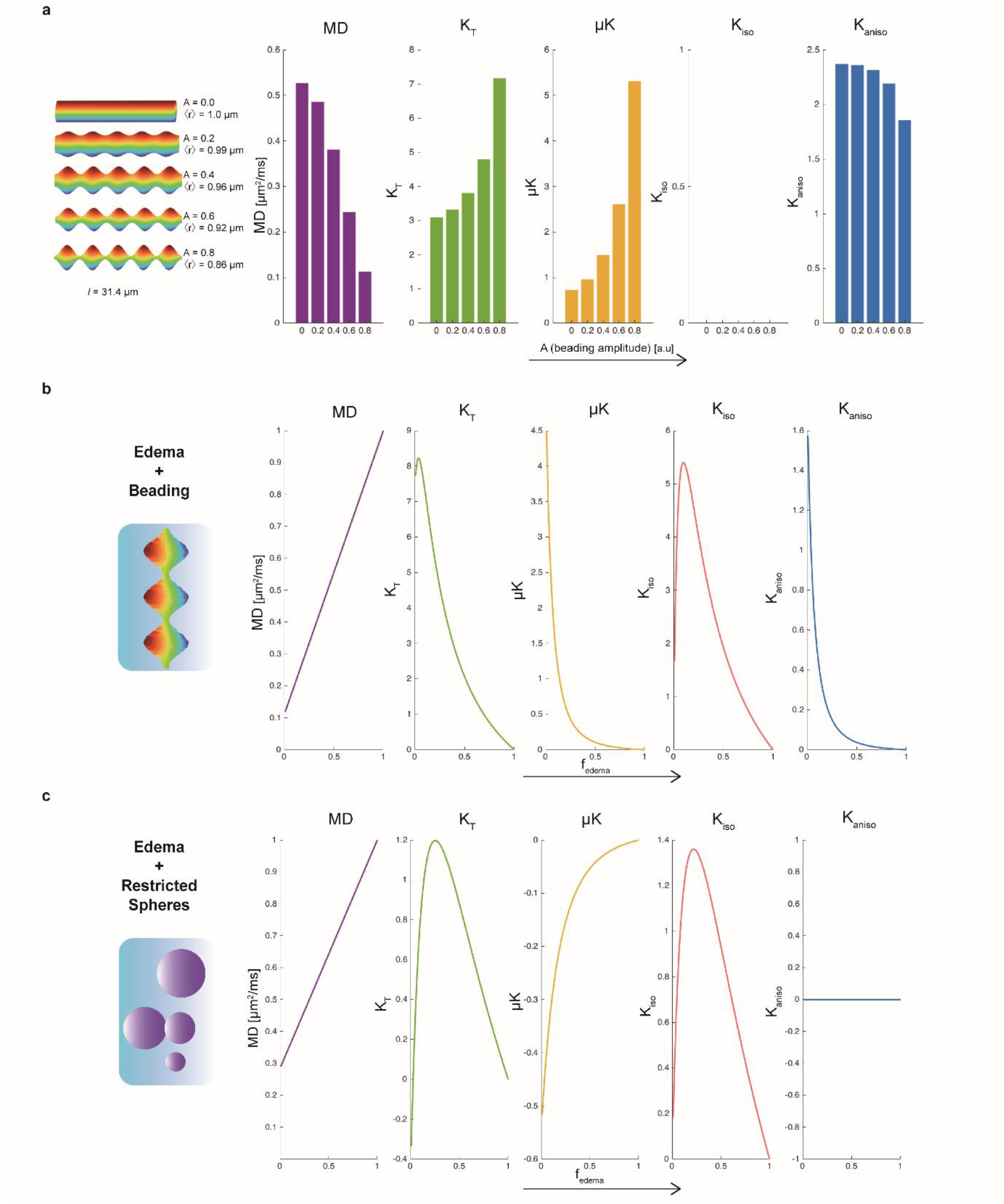
Kurtosis sources estimates with beading and restricted environment (beading and swelling with free water fraction) scenarios. **(a)** Monte Carlo random walk simulations to study the effect of axon beading on the diffusion kurtosis parameters. Five simulated axons were generated by increasing A (which controls the amplitude of beading) from 0, i.e., modulation of axon without beading, to 0.8 (length = 31.4 μm). The respective conventional and novel diffusion kurtosis metrics are presented for all simulated axons scenarios (from left to right: MD [μm^2^/ms], K_T_, μK, K_iso_, K_aniso_). **(b)** Resulting kurtosis estimates from simulated beading effects (A = 0.8) as the free water fraction increases (modulating tissue beading as edema increases). Note that K_iso_ is zero by definition. **(c)** Resulting kurtosis estimates from simulated fully restricted spheres compartments with a generalized Pareto Distribution approximation size distribution (μ= 0.75 ± 0.2 μm) over a range of 100 b-values as free water fraction increases (modulating cell swelling as edema increases).

Interestingly, the introduction of a more rapidly diffusing component into the system (e.g., edema) also explains the stronger WM changes observed in K_iso_: in WM, the microstructure comprises much smaller objects (e.g., axons, myelin etc.) that are characterized by a very low diffusivity. When water with higher diffusivity is introduced to the system, the isotropic variance will be strongly impacted as noted in Fig.6. By contrast, in GM, the initial diffusivity is higher, thereby the variance in diffusivities upon the introduction of a more rapidly diffusing component (e.g., edema) would be smaller than in WM, thereby also making the changes in K_iso_ lower in GM upon ischemia, consistent with our observations.

The anisotropic kurtosis source, K_aniso_ showed a large decrease both in GM and in WM, with much larger decreases in the former. K_aniso_ reflects the microscopic anisotropy without the conflating effects of orientation dispersion suggesting that the spins diffuse in less anisotropic structures upon ischemia, and as shown in Figure 6, a decrease in K_aniso_ would be commensurate with increased beading and edema formation. Thus, the areas shown in Figure 5b very likely reflect the core (areas with large edema and beading effects in the core, while the penumbra is much more subtly affected and shows only small changes in µK, likely reflecting small beading effects due to the ongoing excitotoxicity.

Our findings also have several implications for dMRI modelling in health and disease. First, the often ignored μK clearly needs to be taken into account when modelling dMRI signals in general, as its contribution could clearly be important when looking for changes upon e.g. neurodegeneration. Ignoring μK will result in its contributions becoming conflated into the other kurtosis sources; this in turn could mislead the interpretation of the observed changes in micro-architecture. Finally, it is important to state that characterization of penumbra^50^ – which is a major goal for stroke imaging, could strongly benefit from CTI, as mismatches between the different metrics could be plotted and contrasted with, e.g., changes in perfusion. In addition, the enhanced sensitivity and specificity towards microstructural and edema effects could be used for developing and assessing novel therapies for stroke. More generally, we expect that the CTI metrics will transcend the particular model used here, namely experimental ischemia, and can be generalized towards other important biological mechanisms underlying different diseases, aging, or changes in microstructure associated with development, learning and maturation. All these features bode well for the application of CTI in basic and applied research in the future.

## METHODS

All animal experiments were preapproved by the competent institutional and national authorities and carried out according to European Directive 2010/63.

### Animals

Adult male C57BL/6 J mice (aged 11 weeks, weights 24-29 g, grown with a 12 h /12 h light/dark cycle with ad libitum access to food and water) were used in this study.

### Surgical procedures

A photothrombotic Rose Bengal stroke model^51^ was used to induce a focal infarction in the barrel cortex (S1bf), previously identified upon anatomic mapping, with a solution of Rose Bengal dye (Sigma Aldrich, 95%) dissolved in sterile saline (15 mg/ml) and filtrated through a 0.2 μm sterile filter. Each mouse was injected with meloxicam (Nonsteroidal anti-inflammatory drug) 30 min prior to surgery and anesthetized with isoflurane (∼ 2.5% in 28% oxygen). Temperature was monitored and maintained at 36 – 37°C with a heating pad. The skull was exposed by a median incision of the skin at the dorsal aspect of the head and the periosteum was removed.

The solution was delivered intravenously by a retroorbital injection (10 μl/g animal weight). The animals were then irradiated with a cold light source and a fiber optic light guide (0.89 mm tip) – reaching a colour temperature of 3200 K and a beam light intensity of 10 W/cm^2^ – in the barrel cortex (1.67 mm posterior; 3 mm lateral to bregma^52^) for 15 minutes. A sham group (N = 5) underwent the same conditions except for the lesion-inducing illumination, yet the 15-minute interval post injection was respected.

### Brain extraction and sample preparation

Brain specimens were fixed via transcardial perfusion with 4% Paraformaldehyde (PFA) and extracted from ten adult mice (N = 10) at 3 h post ischemic onset. After extraction from the skull, the brains were immersed in a 4% PFA solution for 24 h and washed in a Phosphate-Buffered Saline solution afterwards, preserved in for at least 24 h. The specimens were subsequently placed in a 10-mm NMR tube filled with Fluorinert (Sigma Aldrich, Lisbon, PT), secured with a stopper to prevent from floating. The tube was then sealed with paraffin film.

### Diffusion MRI acquisition

All *ex vivo* MRI scans were performed on a 16.4 T Aeon Ascend Bruker scanner (Karlsruhe, Germany) equipped with an AVANCE IIIHD console and a Micro5 probe with gradient coils capable of producing up to 3000 mT/m in all directions and a birdcage RF volume coil. Once inserted, the samples were maintained at 37°C due to the probe’s variable temperature capability and were allowed to acclimatize with the surroundings for at least 2 h prior to the beginning of diffusion MRI experiments. In favour of shimming, a B_0_ map was also acquired covering the volume of the brain.

Double diffusion encoding (DDE) data were subsequently acquired for 25 coronal slices using an in house written EPI-based DDE pulse sequence implemented in Paravision 6.0.1 (Bruker BioSpin MRI GmbH, Ettlingen, Germany) and TopSpin 3.1. The diffusion encoding gradient pulse separation Δ and mixing time T_m_ were set to 10 ms, and the pulsed gradient duration δ was set to 1.5 ms. Acquisitions were repeated for five q_1_-q_2_ magnitude combinations (1498 - 0, 1059.2 - 1059.2, 1059.2 - 0, 749.0 - 749.0 and 749.0 - 0 mT/m) corresponding to b-values of 3000 - 0, 1500 - 1500, 1500 - 0, 750 - 750 and 750 - 0 s/mm^2^. Acquisition with q_2_=0 are acquired for 135 directions for q_1_, while equal magnitude q_1_-q_2_ combinations were repeated for 135 parallel and perpendicular directions as described by Henriques et al.^53^. These DDE acquisition-protocol are adjusted according to the requirements for CTI reconstruction^42,53^. In addition, twenty acquisitions without any diffusion-weighted sensitization (zero b-value) were performed for each q_1_-q_2_ magnitude combinations to guarantee a high ratio between the number of non-diffusion and diffusion-weighted acquisitions. For all experiments, the following parameters were used: TR/TE = 3000/49 ms, Field of View = 11 × 11 mm^2^, matrix size 78 × 78, resulting in an in-plane voxel resolution of 141 × 141 μm^2^, slice thickness = 0.5 mm, number of segments = 2, number of averages = 8. For every b_max_ value, the total acquisition time was approximately 1 h and 57 min.

### Structural MRI acquisition

Axial and sagittal T2-weighted images with high resolution and high SNR were acquired for anatomical reference. These data were acquired using RARE pulse sequences with the following parameters: TR = 4000 ms, TE = 50 ms, RARE factor = 12, number of averages = 6. Concerning the axial images, the Field of View (FOV) was 18 × 18 mm^2^, and the matrix size was 240 × 132, resulting in an in-plane voxel resolution of 75 × 76 μm^2^. For the sagittal images, FOV was set to 18 × 9 mm^2^, matrix size to 240 × 120, and subsequently in-plane voxel resolution resulted in 75 × 75 μm^2^ (sagittal). Both axial and sagittal acquisitions sampled the total of 33 slices with a thickness of 0.3 mm. For the coronal images, FOV was established as 10.5 × 10.5 mm^2^, the matrix size as 140 × 140, hence an in-plane voxel resolution of 75 × 75 μm^2^, for 72 slices 0.225 mm thick. The MRI infarct volumes were calculated by the amount of voxels within the designated masks multiplied by the volume of each voxel.

### Diffusion data pre-processing

Before starting the diffusion data analysis, masks for delineation were manually drawn slice-by-slice using Matlab (Matlab R2018b). All datasets were corrected for Gibbs ringing artifacts^54,55^ in Python (Dipy, version 1.0^56^) which suppresses the Gibbs oscillations effects based on sub-voxel Fourier shifts (total variance analysis across three adjacent points for each voxel used to access Gibbs oscillation suppression). All diffusion-weighted datasets underwent realignment via a sub-pixel registration method^57^ in which each set of data for every total diffusion b-value would be realigned to a counter defined dataset with similar DDE gradient pattern combinations.

### CTI reconstruction

CTI was then directly fitted to the data using a linear-least squares fitting procedure implemented to the following equation^42^:

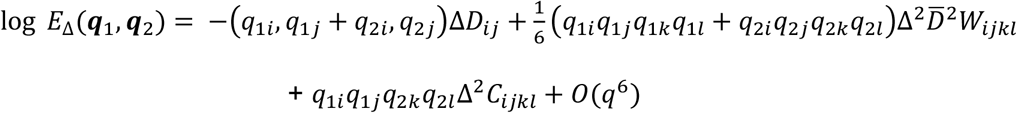

where *D*_*ij*_, *W*_*ijkl*_ and *C*_*ijkl*_ correspond to the diffusion, kurtosis and covariance tensors, respectively. The sources of kurtosis can be extracted from these tensors in the following way^42^: 1) the total kurtosis *K*_*T*_ can be computed from *W*_*ijkl*_; 2) the two inter-compartmental kurtosis sources (*K*_*aniso*_ and *K*_*iso*_) can be extracted from *C*_*ijkl*_; and 3) an intra-compartmental kurtosis source *K*_*intra*_ can be estimated as *K*_*T*_ − *K*_*aniso*_ − *K*_*iso*_.

### Region of Interest Analysis

A region of Interest (ROI) analysis was performed by manual selection of the most relevant areas for every estimate data (MD, K_T_, μK, K_iso_ and K_aniso_) slice in both hemispheres in all brains, containing the total lesion and counterpart region in the opposite hemisphere, based on interhemispheric visual asymmetry (Supplementary Fig. S5). Within these more general ROIs, GM and WM regions were further selected according to a manual selection of WM regions based on an anatomical brain map^52^ and FA measures. All ROI analyses were performed in Matlab (Matlab R2018b).

### Interhemispheric ratio analysis

An interhemispheric ratio analysis was performed by calculating interhemispheric ratios percentage changes between ipsilesional and contralesional (stroked brain) hemispheres (N = 5) as presented below:

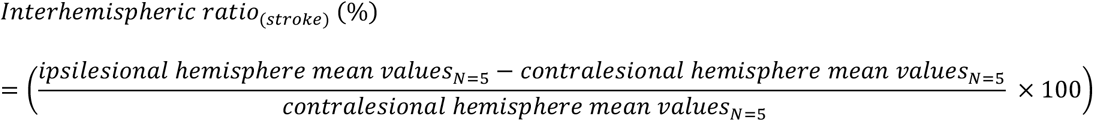

### Statistical analysis features

A Shapiro-Wilk test was used to test normality (p < 0.05) over the hemispheric ratio data for all affected voxels, GM and WM. Upon normality validation for total affected voxels and GM data, a one-way ANOVA test (p < 0.05) was performed to test if the interhemispheric mean differences were significant between the different metrics (MD, K_T_, μK, K_iso_ and K_aniso_) in these regions, with a subsequent Tukey’s HSD multiple comparison test to assess which metrics means were statistically significant (p < 0.05). A non-parametric Kruskal-Wallis test (p < 0.05) was used to test if the interhemispheric mean differences were significant between the same metrics in WM, followed by the Dunn’s multiple comparison test to assess the statistically different ratios (p < 0.05).

All statistical analysis were performed in Matlab (Matlab R2018b).

### Data visualization

All volumetric datasets were rendered with ImageJ. Volume render maps were produced using the plugin Volume Viewer^58^, with tricubic smooth interpolation for K_T_ and μK sources with the following settings: threshold = 0.5 – 1.5, “display_mode=4”, “axes=0”, “interpolation=2”, “useLight=0”, “specularValue=0.5”, “diffuseValue=0.5”, “shineValue=0”, “ambientValue=0”, “objectLightValue=2”, “bg_r=255”, “bg_g=255”, “bg_b=255” and “lut=3” (7 frame per second volumetric video renders are displayed in Supplementary Videos). MRIcroGL^59^ was used to create 2D lesion overlay maps on the averaged b0-weighted images. NIFTI format images from each metric affected voxels ROI (GM + WM) and respective estimates were overlayed with the following settings: MD coloured in “2green”, scale: 0.2 – 0.7, opacity: 100%; μK coloured in “7cool”, scale: 0 – 1, opacity = 70%; K_T_ coloured in “6warm”, scale: 0 – 2, opacity = 100%; K_iso_ coloured in “8redyell”, scale: 0 – 10, opacity = 100%; K_aniso_ coloured in “Plasma”, scale: 0 – 2, opacity = 100%;

### Simulations

Monte Carlo random walk simulations were performed to study the effect of axon beading on the diffusion kurtosis parameters. An axon without beading was modelled as an impermeable cylinder (length = 31.4 μm, radius = 1 μm) aligned with the z-axis. Beading was introduced by making the radius depend on the location along the z-axis: r(z) = r_0_ + A · sin(B · z), where r_0_ is the mean radius, A ∈[0,1] controls the amplitude of beading, and B controls the frequency and location of beading. Five simulated axons, shown in Figure X, were generated by increasing A from 0 to 0.8 in equal steps and adjusting r_0_ to keep the volume constant. B = 1 μm^-1^ for all simulated axons (Fig. 6a).

Simulated data was generated using Disimpy^60^ with 10^5^ random walkers, 10^4^ time steps, and diffusivity of 2 μm^2^/ms. The initial positions of the random walkers were randomly sampled from a uniform distribution inside the axons. MR signals were generated using pulsed gradient single diffusion encoding with δ = 1.5 ms and Δ = 10 ms, 60 diffusion encoding directions uniformly distributed over the surface of half a sphere, and 10 b-values uniformly distributed between 0 and 3 ms/μm^2^. The diffusion and kurtosis tensors were estimated from the simulated signals using a weighted linear least squares fit in Dipy^56^. MD, μK, K_iso_, and K_aniso_ were calculated from the diffusion tensor eigenvalues and the elements of the kurtosis tensor^42^ for individual components (Fig. 6a).

Analytically computed pulsed gradient spin echo sequence signals were simulated for two different models using the MISST toolbox^61–64^: beading (to model beaded axons) with formation of edema, assuming the most beaded previously simulated scenario with *A* = 0.8 (Fig. 6b); spherical compartments with restricted diffusion (to model cell swelling) and formation of edema, with combinations of radii between 0.8 μm and 10 μm, with a diffusivity value of 2 μm^2^/ms (Fig. 6c).

### Nissl Cresyl Violet staining

Histological analysis was performed in one of the stroked *ex vivo* samples. Slices were obtained through Vibratome sectioning with a thickness of 0.04 mm and Mowiol containing 2.5 % 1,4 diazobicyclo-[2.2.2]-octane (DABCO, Sigma, D2522) was used as the mounting media. The brain sections were then fixed with 10% formalin and processed with Nissl-Cresyl Violet staining for microscopy in order to assess tissue damage and cell loss within the infarcted region.

### Optical imaging

Histological imaging was performed with a ZEISS Axio Scan.Z1 (Zeiss, Germany) coupled to a Hitachi 3 CCD colour camera and processed with QuPath 0.2.3 (Fig. 2e). Images were magnified (53X) for both ipsi- and contralesional hemispheres.

### Animal monitoring for *in vivo* MRI imaging

For anaesthesia induction, the body temperature of the mouse was kept constant by placing the animal on top of an electrical heating pad. Anaesthesia with a mixture of medical air and 4% isoflurane (Vetflurane, Virbac, France) was maintained until the animal righting reflex and any reaction to firm foot squeeze were lost. The isoflurane concentration was regulated and reduced to 2.5%. The mouse was then weighed and transferred to the animal bed, prone positioned above a heated water pad – in order for the mouse body temperature not to oscillate during the experiments –, having its head placed with its upper incisors held on to a mouth bite bar. Oxygen concentrations were kept between 27% and 28%, monitored by a portable oxygen monitor (MX300-I, Viamed, United Kingdom). Ear bars were used for a safe and efficient head fixation (into external meatus) and eye ointment (Bepanthen Eye Drops, Bepanthen, Germany) was applied to prevent the corneas from drying. A rectal temperature probe and a respiration sensor (Model 1030 Monitoring Gating System, SAII, United States of America) were placed for real-time monitoring of these physiological measurements to guarantee the animal’s welfare and immobilization. Considering the water molecules sensitivity towards temperature alterations, the waterbed temperature was cautiously monitored and controlled to avoid oscillations. Respiration rates were also monitored and maintained at physiological levels throughout dMRI scanning.

### *in vivo* MRI acquisitions

The *in vivo* MRI data were acquired on a 9.4 T horizontal MRI scanner (BioSpec 94/20 USR, Bruker BioSpin, Germany) equipped a gradient system able to produce up to 660 mT/m in every direction, an 86 mm quadrature coil for transmission and a 4-element array surface cryocoil for reception.

Sagittal T2-weighted images were acquired for anatomical reference using a RARE pulse sequence with the following parameters: TR = 2000 ms, TE = 36 ms, RARE factor = 8, number of averages = 8. The field of view was 24 × 16.1 mm^2^, the matrix size was 256 × 256, resulting in an in-plane voxel resolution of 150 × 150 μm^2^. The slice thickness was 0.5 mm, and 21 slices were sampled.

Following an optimized protocol when compared to the *ex vivo* experiment, DDE data were acquired for 5 coronal slices using our in house written EPI-based DDE pulse sequence. The diffusion encoding gradient pulse separation Δ and mixing time τ_m_ were set to 10 ms, and the pulsed gradient duration δ was set to 4 ms. Acquisitions were repeated for five q_1_-q_2_ magnitude combinations (518.79 - 0, 366.84 – 366.84.2, 366.84 - 0, 259.4 – 259.4 and 259.4 - 0 mT/m) corresponding to b-values of 3000 - 0, 1500 - 1500, 1500 - 0, 750 - 750 and 750 - 0 s/mm^2^. For each gradient combination, experiments are repeated for the same directions in the *ex vivo* experiments and described by Henriques et al.^53^. In addition, twenty acquisitions without any diffusion-weighted sensitization (b-value = 0) were performed. For all experiments, the following parameters were used: TR/TE = 2800/44.5 ms, FOV = 12 × 12 mm^2^, matrix size 78 × 78, resulting in an in-plane voxel resolution of 181 × 181 μm^2^, slice thickness = 0.85 mm, number of segments = 1, number of averages = 1 and partial Fourier acceleration of 1.25. For every b_max_ value, the total acquisition time was approximately 7 min.

## Data availability

The data sets generated and analysed during the current study are available from the corresponding author upon reasonable request.

## Code availability

Custom MATLAB code for dMRI pre- and post-processing of data is available from the corresponding author upon reasonable request.

## ACKNOWLEDGMENTS

This study was funded by the European Research Council (ERC) under the European Union’s Horizon 2020 research and innovation programme (Starting Grant, agreement No. 679058). The authors acknowledge the vivarium of the Champalimaud Centre for the Unknow, a facility of CONGENTO financed by Lisboa Regional Operational Programme (Lisboa 2020), project LISBOA01-0145-FEDER-022170, and also the Champalimaud Histopathology and the Champalimaud ABBE Platforms. The authors also want to thank Ms. Beatriz Cardoso for assistance in the preparation of the *ex vivo* mouse brain specimens.

## REFERENCES

1. Suter, T.A.C.S. & Jaworski, A. Cell migration and axon guidance at the border between central and peripheral nervous system. Science. 365, 881–891 (2019).

2. Hughes, E. G., Orthmann-Murphy, J. L., Langseth, A. J. & Bergles, D. E. Myelin remodeling through experience-dependent oligodendrogenesis in the adult somatosensory cortex. Nat. Neurosci. 21, 696–706 (2018).

3. Brodt, S. et al. Fast track to the neocortex: A memory engram in the posterior parietal cortex. Science. 362, 1045–1048 (2018).

4. Zatorre, R. J., Fields, R. D. & Johansen-Berg, H. Plasticity in gray and white: neuroimaging changes in brain structure during learning. Nat. Neurosci. 15, 528–536 (2012).

5. Scholz, J., Klein, M. C., Behrens, T. E. J. & Johansen-Berg, H. Training induces changes in white-matter architecture. Nat. Neurosci. 12, 1370–1371 (2009).

6. Craddock, R. C. et al. Imaging human connectomes at the macroscale. Nat. Methods 10, 524–539 (2013).

7. Pestilli, F., Yeatman, J. D., Rokem, A., Kay, K. N. & Wandell, B. A. Evaluation and statistical inference for human connectomes. Nat. Methods 11, 1058–1063 (2014).

8. Hill, R. A., Li, A. M. & Grutzendler, J. Lifelong cortical myelin plasticity and age-related degeneration in the live mammalian brain. Nat. Neurosci. 21, 683–695 (2018).

9. Murphy, T. H. & Corbett, D. Plasticity during stroke recovery: From synapse to behaviour. Nat. Rev. Neurosci. 10, 861–872 (2009).

10. Li, N. et al. mTOR-Dependent Synapse Formation Underlies the Rapid Antidepressant Effects of NMDA Antagonists. Science. 329, 959–964 (2010).

11. Peelaerts, W. et al. α-Synuclein strains cause distinct synucleinopathies after local and systemic administration. Nature 522, 340–344 (2015).

12. De Strooper, B. & Karran, E. The Cellular Phase of Alzheimer’s Disease. Cell 164, 603– 615 (2016).

13. Budde, M. D., Janes, L., Gold, E., Turtzo, L. C. & Frank, J. A. The contribution of gliosis to diffusion tensor anisotropy and tractography following traumatic brain injury: validation in the rat using Fourier analysis of stained tissue sections. Brain 134, 2248– 2260 (2011).

14. Mathers, C. et al. WHO methods and data sources for country-level causes of death 2000-2015. (2017). Available at: https://www.who.int/healthinfo/global_burden_disease/GlobalCOD_method_2000_2015.pdf?ua=1.

15. Belov Kirdajova, D., Kriska, J., Tureckova, J. & Anderova, M. Ischemia-Triggered Glutamate Excitotoxicity From the Perspective of Glial Cells. Front. Cell. Neurosci. 14, (2020).

16. Budde, M. D. & Frank, J. A. Neurite beading is sufficient to decrease the apparent diffusion coefficient after ischemic stroke. Proc. Natl. Acad. Sci. U. S. A. 107, 14472– 14477 (2010).

17. Skinner, N. P., Kurpad, S. N., Schmit, B. D. & Budde, M. D. Detection of acute nervous system injury with advanced diffusion-weighted MRI: a simulation and sensitivity analysis. NMR Biomed. 28, 1489–1506 (2015).

18. Zeiler, S. R. et al. Paradoxical Motor Recovery From a First Stroke After Induction of a Second Stroke. Neurorehabil. Neural Repair 30, 794–800 (2016).

19. Veraart, J. et al. Noninvasive quantification of axon radii using diffusion MRI. Elife 9, (2020).

20. Andersson, M. et al. Axon morphology is modulated by the local environment and impacts the noninvasive investigation of its structure–function relationship. Proc. Natl. Acad. Sci. 117, 33649–33659 (2020).

21. Novikov, D. S., Jensen, J. H., Helpern, J. A. & Fieremans, E. Revealing mesoscopic structural universality with diffusion. Proc. Natl. Acad. Sci. 111, 5088–5093 (2014).

22. Novikov, D. S., Fieremans, E., Jensen, J. H. & Helpern, J. A. Random walks with barriers. Nat. Phys. 7, 508–514 (2011).

23. Le Bihan, D. Looking into the functional architecture of the brain with diffusion MRI. Nat. Rev. Neurosci. 4, 469–480 (2003).

24. Moseley, M. E. et al. Early detection of regional cerebral ischemia in cats: Comparison of diffusion- and T2-weighted MRI and spectroscopy. Magn. Reson. Med. 14, 330–346 (1990).

25. Karbasforoushan, H., Cohen-Adad, J. & Dewald, J. P. A. Brainstem and spinal cord MRI identifies altered sensorimotor pathways post-stroke. Nat. Commun. 10, 3524 (2019).

26. Basser, P. J., Mattiello, J. & LeBihan, D. MR diffusion tensor spectroscopy and imaging. Biophys. J. 66, 259–267 (1994).

27. Jensen, J. H., Helpern, J. A., Ramani, A., Lu, H. & Kaczynski, K. Diffusional kurtosis imaging: The quantification of non-gaussian water diffusion by means of magnetic resonance imaging. Magn. Reson. Med. 53, 1432–1440 (2005).

28. Yin, J. et al. Diffusion Kurtosis Imaging of Acute Infarction: Comparison with Routine Diffusion and Follow-up MR Imaging. Radiology 287, 651–657 (2018).

29. Muir, K. W., Buchan, A., von Kummer, R., Rother, J. & Baron, J.-C. Imaging of acute stroke. Lancet Neurol. 5, 755–768 (2006).

30. Stejskal, E. O. & Tanner, J. E. Spin diffusion measurements: Spin echoes in the presence of a time-dependent field gradient. J. Chem. Phys. 42, 288–292 (1965).

31. Shemesh, N. et al. Conventions and Nomenclature for Double Diffusion Encoding NMR and MRI. Magn. Reson. Med. 87, 82–87 (2016).

32. Lee, H. H., Papaioannou, A., Kim, S. L., Novikov, D. S. & Fieremans, E. A time-dependent diffusion MRI signature of axon caliber variations and beading. Commun. Biol. 3, (2020).

33. Xing, C., Arai, K., Lo, E. H. & Hommel, M. Pathophysiologic cascades in ischemic stroke. Int. J. Stroke 7, 378–385 (2012).

34. Stokum, J. A., Gerzanich, V. & Simard, J. M. Molecular pathophysiology of cerebral edema. J. Cereb. Blood Flow Metab. 36, 513–538 (2016).

35. Baron, C. A. et al. Reduction of Diffusion-Weighted Imaging Contrast of Acute Ischemic Stroke at Short Diffusion Times. Stroke 46, 2136–2141 (2015).

36. Novikov, D. S., Fieremans, E., Jespersen, S. N. & Kiselev, V. G. Quantifying brain microstructure with diffusion MRI: Theory and parameter estimation. NMR Biomed. 32, e3998 (2019).

37. Szafer, A., Zhong, J. & Gore, J. C. Theoretical Model for Water Diffusion in Tissues. Magn. Reson. Med. 33, 697–712 (1995).

38. Weber, R. A. et al. Diffusional kurtosis and diffusion tensor imaging reveal different time-sensitive stroke-induced microstructural changes. Stroke 46, 545–550 (2015).

39. Nørhøj, S. & Buhl, N. The displacement correlation tensor?: Microstructure, ensemble anisotropy and curving fibers. J. Magn. Reson. Imaging 208, 34–43 (2011).

40. Mitra, P. P. Multiple wave-vector extensions of the NMR pulsed-field-gradient spin-echo diffusion measurement. Phys. Rev. 51, 74–78 (1995).

41. Jespersen, S. N., Lundell, H., Sønderby, C. K. & Dyrby, T. B. Orientationally invariant metrics of apparent compartment eccentricity from double pulsed field gradient diffusion experiments. NMR Biomed. 26, 1647–1662 (2013).

42. Henriques, R. N., Jespersen, S. N. & Shemesh, N. Correlation tensor magnetic resonance imaging. Neuroimage 211, 116605 (2020).

43. Kerkelä, L., Henriques, R. N., Hall, M. G., Clark, C. A. & Shemesh, N. Validation and noise robustness assessment of microscopic anisotropy estimation with clinically feasible double diffusion encoding MRI. Magn. Reson. Med. 1–13 (2019).

44. Nilsson, M. et al. Tensor-valued diffusion MRI in under 3 minutes: an initial survey of microscopic anisotropy and tissue heterogeneity in intracranial tumors. Magn. Reson. Med. 83, 608–620 (2020).

45. Callaghan, P. T., Coy, A., MacGowan, D., Packer, K. J. & Zelaya, F. O. Diffraction-like effects in NMR diffusion studies of fluids in porous solids. Nature 351, 467–469 (1991).

46. Novikov, D. S. & Kiselev, V. G. Effective medium theory of a diffusion-weighted signal. NMR Biomed. 23, 682–697 (2010).

47. Hui, E. S. et al. Stroke assessment with diffusional kurtosis imaging. Stroke 43, 2968– 2973 (2012).

48. Umesh Rudrapatna, S. et al. Can diffusion kurtosis imaging improve the sensitivity and specificity of detecting microstructural alterations in brain tissue chronically after experimental stroke? Comparisons with diffusion tensor imaging and histology. Neuroimage 97, 363–373 (2014).

49. Takeuchi, H. et al. Neuritic Beading Induced by Activated Microglia Is an Early Feature of Neuronal Dysfunction Toward Neuronal Death by Inhibition of Mitochondrial Respiration and Axonal Transport. J. Biol. Chem. 280, 10444–10454 (2005).

50. Astrup, J., Siesjö, B.K. & Symon, L. Thresholds in cerebral ischemia -the ischemic penumbra. Stroke 12, 723–725 (1981).

51. Watson, B. D., Dietrich, W. D., Busto, R., Wachtel, M. S. & Ginsberg, M. D. Induction of reproducible brain infarction by photochemically initiated thrombosis. Ann. Neurol. 17, 497–504 (1985).

52. Franklin, K. & Paxinos, G. Paxinos and Franklin’s the Mouse Brain in Stereotaxic Coordinates. (2019).

53. Henriques, R. N., Jespersen, S. N. & Shemesh, N. Evidence for microscopic kurtosis in neural tissue revealed by Correlation Tensor MRI. (2021). 2102.11701

54. Henriques, R. N. Advanced Methods for Diffusion MRI Data Analysis and their Application to the Healthy Ageing Brain. (Cambridge University, 2018).

55. Kellner, E., Dhital, B., Kiselev, V. G. & Reisert, M. Gibbs-ringing artifact removal based on local subvoxel-shifts. Magn. Reson. Med. 76, 1574–1581 (2016).

56. Garyfallidis, E. et al. Dipy, a library for the analysis of diffusion MRI data. Front. Neuroinform. 8, (2014).

57. Guizar-sicairos, M., Thurman, S. T. & Fienup, J. R. Guizar_Efficent_subpixel_. Opt. Lett. 33, 156–158 (2008).

58. Barthel, K. U. Volume Viewer. (2012). Available at: https://imagej.net/plugins/volume-viewer.html.

59. Rorden, C. & Brett, M. Stereotaxic display of brain lesions. Behav. Neurol. 12, 191–200 (2000).

60. Kerkelä, L., Nery, F., Hall, M. & Clark, C. Disimpy: A massively parallel Monte Carlo simulator for generating diffusion-weighted MRI data in Python. J. Open Source Softw. 5, 2527 (2020).

61. Drobnjak, I., Siow, B. & Alexander, D. C. Optimizing gradient waveforms for microstructure sensitivity in diffusion-weighted MR. J. Magn. Reson. 206, 41–51 (2010).

62. Drobnjak, I., Zhang, H., Hall, M. G. & Alexander, D. C. The matrix formalism for generalised gradients with time-varying orientation in diffusion NMR. J. Magn. Reson. 210, 151–157 (2011).

63. Ianuş, A., Siow, B., Drobnjak, I., Zhang, H. & Alexander, D. C. Gaussian phase distribution approximations for oscillating gradient spin echo diffusion MRI. J. Magn. Reson. 227, 25–34 (2013).

64. Ianuş, A., Alexander, D. C. & Drobnjak, I. Microstructure Imaging Sequence Simulation Toolbox. in 34–44 (2016).

