## Supplementary Figure for "Unraveling micro-architectural modulations in neural tissue upon ischemia by Correlation Tensor MRI"

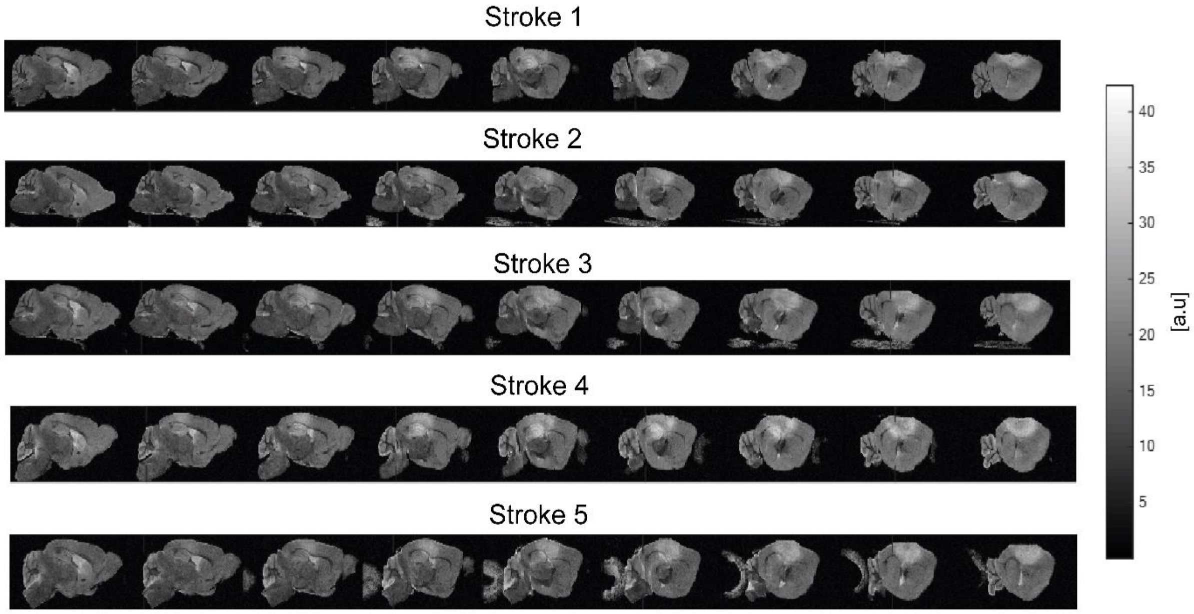


**Supplementary Figure S1 –** Raw T2-weighted images from all ex vivo brains 3 h post stroke onset for reproducibility analysis. 9 representative sagittal slices are presented from right to left. The ischemic territory is delimited by the hyperintense voxels (brighter subcortical regions in the somatosensory cortex).

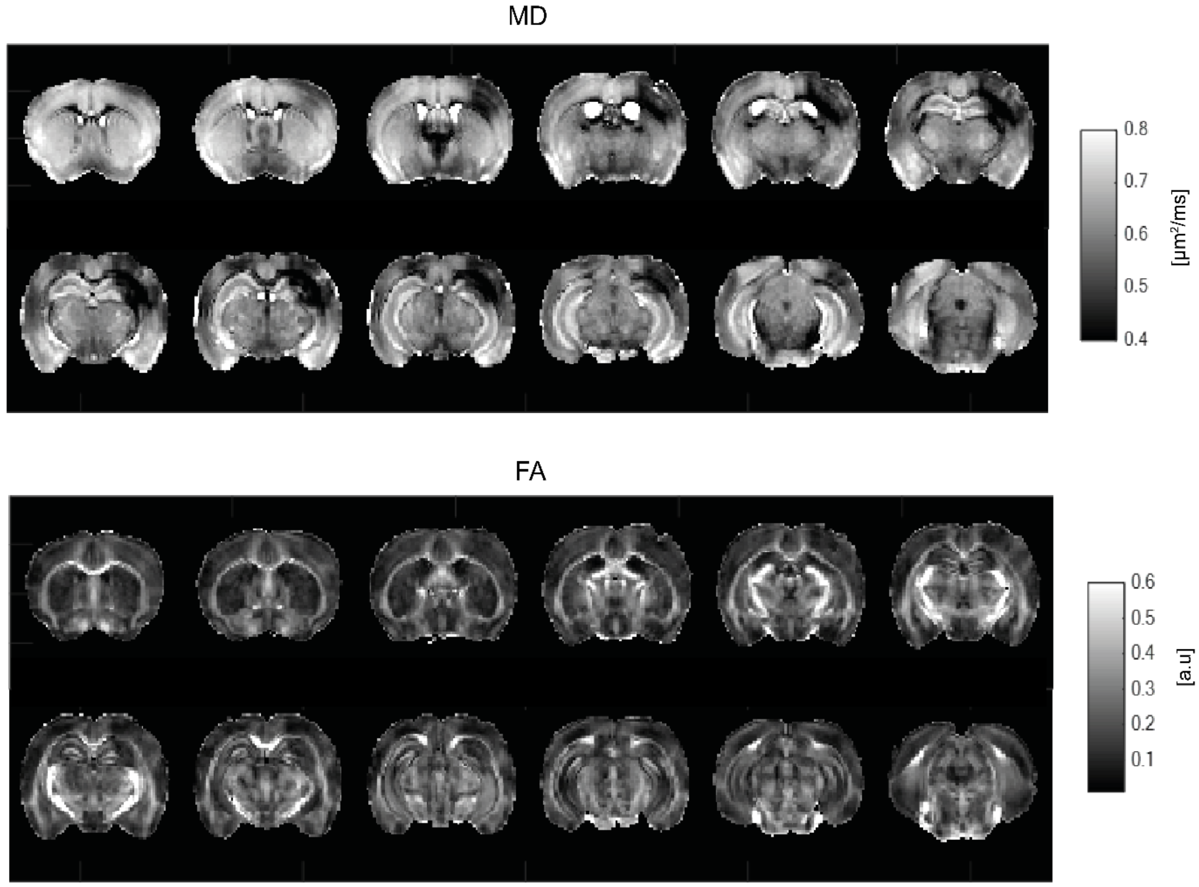


**Supplementary Figure S2 –** Mean Diffusivity (MD) ($\mu m^2/ms$) and Fractional Anisotropy (FA) maps for 12 slices from a representative stroked brain.

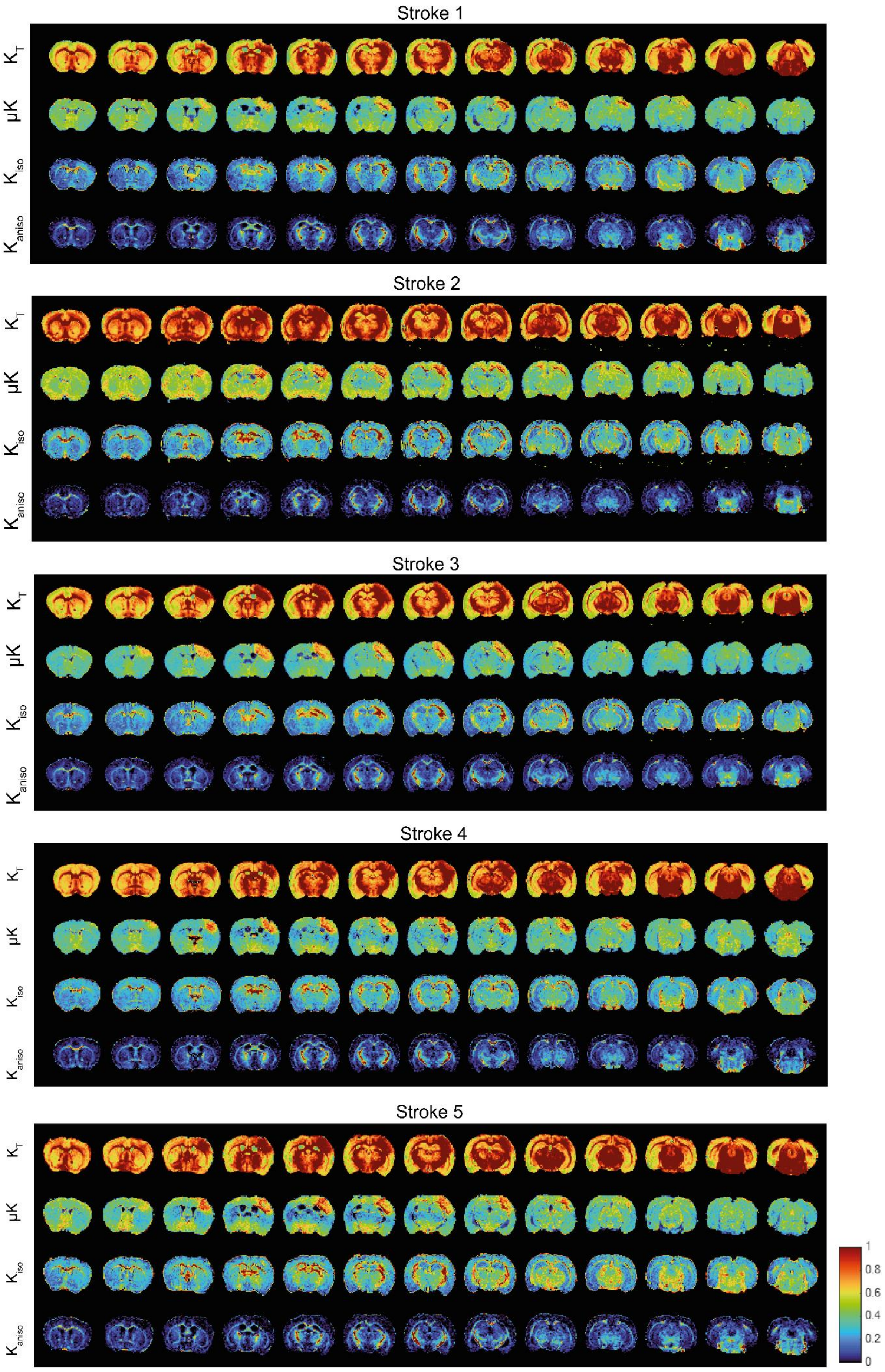


**Supplementary Figure S3 –** Reproducibility of CTI estimates ($K_T$, μK, $K_{iso}$ and $K_{aniso}$) over all 5 stroked mouse brains.

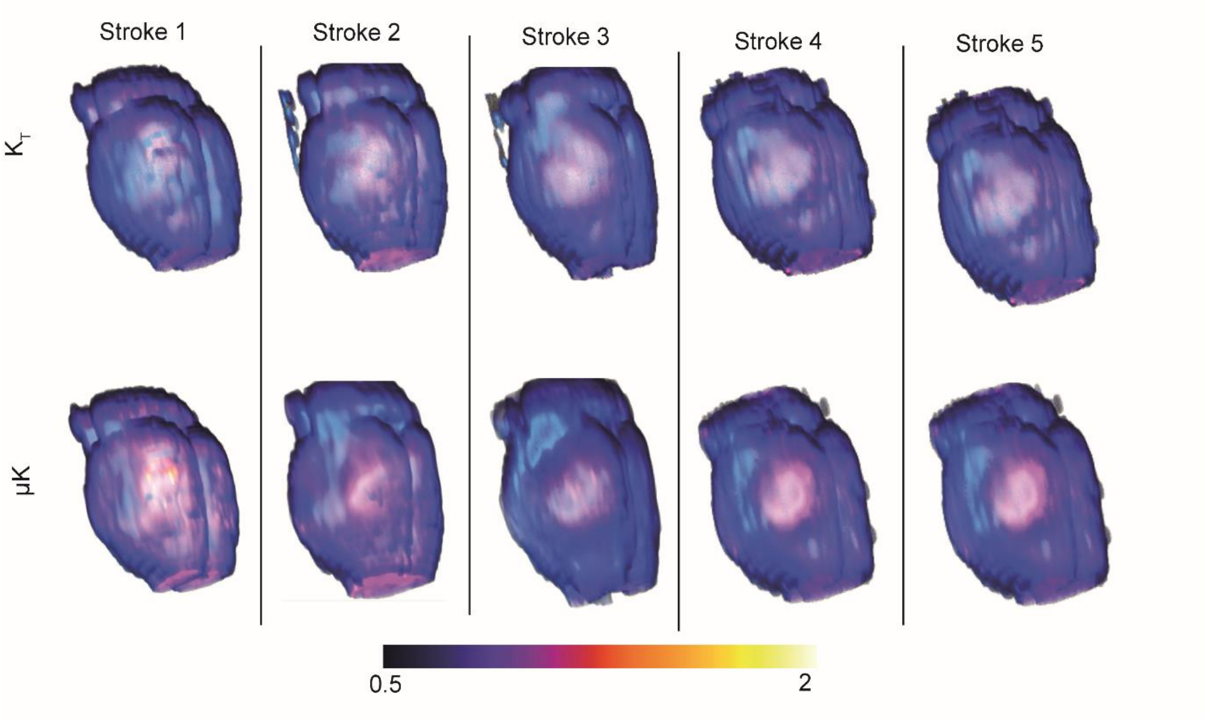


**Supplementary Figure S4 –** 3D render of $K_T$ and μK from all five stroked brains using tricubic smooth interpolation for $K_T$ and μK sources estimates.

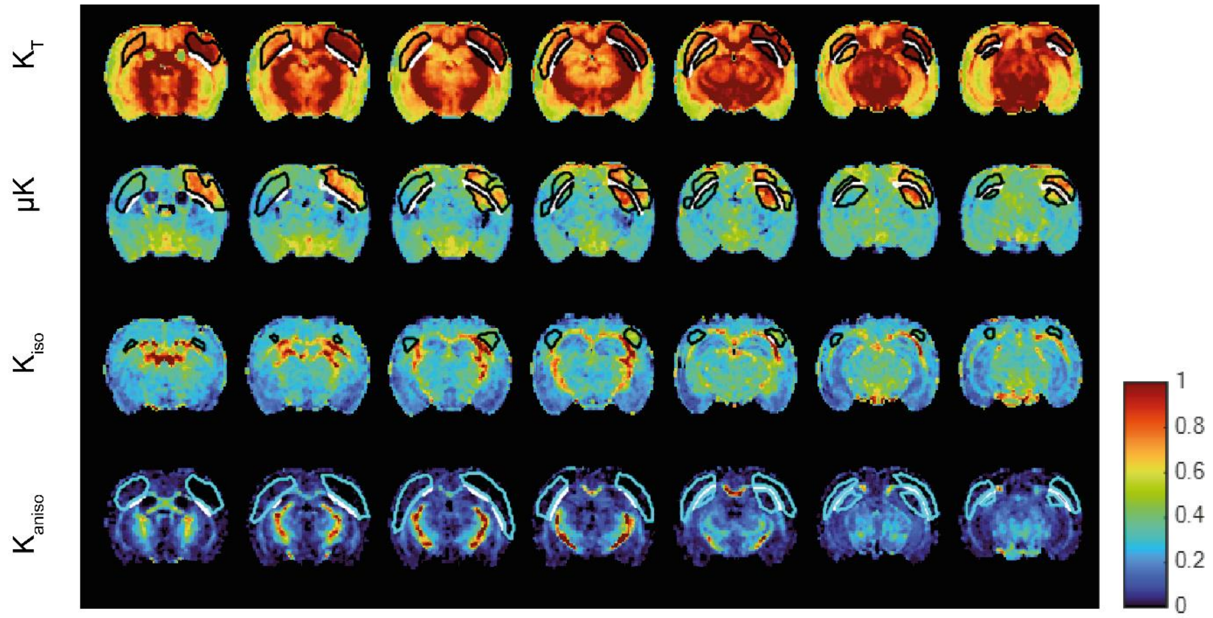


**Supplementary Figure S5 –** Seven representative slices from $K_T$, μK, $K_{iso}$ and $K_{aniso}$ maps of an extracted brain from the stroke group presenting manually defined ROIs. GM ROIs are delineated in black for $K_T$, μK and $K_{iso}$; in cyan for $K_{aniso}$, whereas WM ROIs are delineated in white. Total lesion ROIs (total affected voxels) correspond to the sum of the area covered by GM and WM ROIs. Total ROIs were manually drawn upon qualitative analysis of the interhemispheric asymmetry and WM ROIs were manually drawn based on FA maps and anatomical reference.

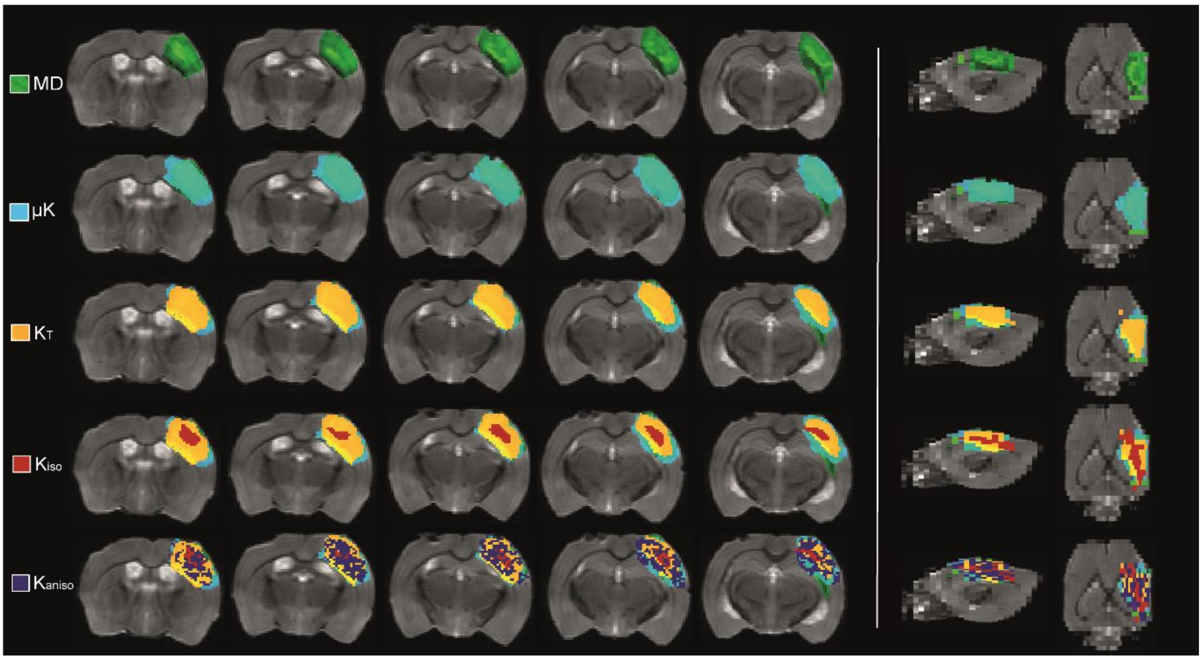


**Supplementary Figure S6 –** Metrics addition analysis using MRIcroGL in zero b-value averaged images of a representative stroked brain (stroke 5). 2D renderings of overlay ROI maps on the ipsilesional hemisphere were performed according to the voxels affected by the lesion in each metric, with the following order: MD (green), μK (cyan), $K_T$ (yellow), $K_{iso}$ (red) and $K_{aniso}$ (dark purple). Sagittal and axial views are presented on the right panel, respectively.

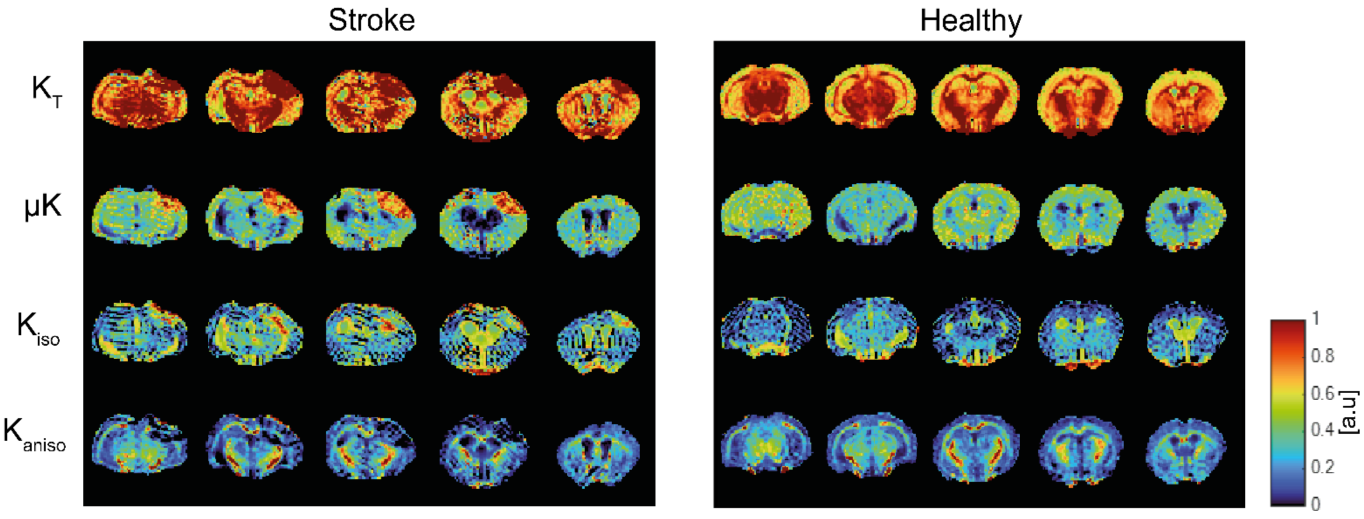


**Supplementary Figure S7 –** Kurtosis sources map for *in vivo* acquisitions from a stroked mouse and healthy mouse. In the stroked animal, the $K_T$ estimate presents high intensity values in the lesion area, consistent with the *ex vivo* maps. μK reveals hyperintense values in the lesioned region in the ipsilesional hemisphere, for both white and gray matter. $K_{iso}$ shows elevated intensity values in white matter and $K_{aniso}$ has low values within the lesioned region (both white and gray matter). In the healthy animal, $K_T$ presents higher values than any other sources. $K_{aniso}$ values reveal high intensities in white matter, whereas $K_{iso}$ shows lower intensity values for both white and gray matter, consistent with the *ex vivo* control brains.
